# Field evaluation of malachite green loop-mediated isothermal amplification as a malaria parasite detection tool in a health post in Roraima state, Brazil

**DOI:** 10.1101/408609

**Authors:** Heather M. Kudyba, Jaime Louzada, Dragan Ljolje, Karl A. Kudyba, Vasant Muralidharan, Joseli Oliveira-Ferreira, Naomi W. Lucchi

## Abstract

Malaria is a debilitating parasitic disease that causes significant morbidity and mortality. Microscopic detection of parasites is currently the “gold standard” diagnostic. This technique is limited in its ability to detect low-density infections, is time consuming, and requires a highly trained microscopist. Malaria epidemiological surveillance studies especially aimed at the detection of low-density infection and asymptomatic cases will require more sensitive and user-friendly tools. We have shown previously that the molecular-based, colorimetric malachite green loop-mediated isothermal amplification (MG-LAMP) assay is a valuable tool for diagnosing malaria infection in a laboratory setting. In this study, we field evaluated this assay in a malaria diagnostic post in Roraima, Brazil. We prospectively collected 91 patient samples and performed microscopy, MG-LAMP, and real-time PCR (PET-PCR) to detect *Plasmodium* infection. Two independent readers were used to score the MG-LAMP tests to assess whether the sample was positive (blue/green) or negative (clear). There was 100% agreement between the two readers (Kappa=1). All tests detected 33 positive samples, but both the MG-LAMP and PET-PCR detected 6 and 7 more positive samples, respectively. The PET-PCR assay detected 6 mixed infections (defined as infection with both *P. falciparum* and *P. vivax*) while microscopy detected one and MG-LAMP detected two of these mixed infections. Microscopy did not detect any *Plasmodium* infection in 26 of the enrolled asymptomatic cases while MG-LAMP detected five and PET-PCR assay three positive cases. Overall, MG-LAMP provided a simpler and user-friendly molecular method for malaria diagnosis that is more sensitive than microscopy. Additionally, MG- LAMP has the capacity to test 38 samples per run (one hour), allowing for the screening of large number of samples which is appealing when large-scale studies are necessary e.g. in community surveillance studies. The current MG-LAMP assay was limited in its ability to detect mixed infection when compared to the PET-PCR, but otherwise proved to be a powerful tool for malaria parasite detection in the field and opens new perspectives in the implementation of surveillance studies in malaria elimination campaigns.

## INTRODUCTION

Malaria is a devastating disease that remains a major global health burden. This illness arises from infection with parasites of the genus *Plasmodium*. Cases of the most significant morbidity and mortality in humans are caused by the most prevalent species, *P. vivax* and *P. falciparum*. *P. ovale* and *P. malariae* also cause human malaria, but the infections are typically associated with milder symptoms. The treatment regimens given to patients infected with different species of *Plasmodium* may vary, thus accurate diagnosis is imperative^1^. Currently, the primary method used in Brazil for the diagnosis of *Plasmodium* is microscopy of a Giemsa-stained thick or thin blood smear. While this technique is the gold standard for malaria diagnosis due to low cost and accessibility, it has limitations, including the inability to detect parasites when the parasitemia is extremely low, occasional misdiagnosis of mixed-species infection, and this technique is extremely time consuming^2^-^6^. Rapid diagnostic test (RDT’s) provide their own set of limitations including the presence of circulating parasites carrying deletions of genes encoding the antigens being detected by the test and interpretation issues with low density infections^7^-^12^. Malaria elimination efforts are hampered by the lack of sensitive tools to detect infections with low-level parasitemia, usually below the threshold of standard diagnostic methods, microscopy and rapid diagnostic tests. The elimination of malaria will require active case detection in low transmission areas as well as the ability to detect sub-microscopic infections^13^. This necessitates a diagnostic tool that can test many samples at once and detect malaria in patients with low parasite densities. Thus, the implementation of more sensitive diagnostic tools in the field is of the utmost importance.

Molecular-based diagnostic tools provide more sensitive and specific methods for detecting *Plasmodium* infections than microscopy and RDTs. To be a “significant improvement” over expert microscopy, it is recommended that molecular tests be at least one log more sensitive than microscopy; therefore, have a detection limit of 2 parasites/µl or less (WHO, 2014). The use of molecular-based diagnostic tools in research and in epidemiological surveys has expanded in recent years. However, their use is still limited to laboratories with more sophisticated facilities due to the requirement of specialized equipment and technical expertise. Simpler molecular tests, such as the loop-mediated isothermal amplification (LAMP) assays, promise to facilitate the use of molecular tests even in facilities with limited resources^14^-^18^. Recently, we reported on the feasibility and sensitivity of a *Plasmodium* genus-specific malachite green loop-mediated isothermal amplification (MG-LAMP) as a method for diagnosing *Plasmodium* infection^19^. The MG-LAMP is a colorimetric LAMP assay that relies on the visual readout of the results as positive (green color) or negative (colorless) post amplification at a constant temperature. Amplification is performed in a 40-well mini heat-block, allowing for many samples to be ran at once.

In this study, we field-tested the practicality and effectiveness of this tool in Roraima, Brazil using freshly isolated patient samples in local health clinics. Previously reported *P.* falciparum^20^ and *P. vivax*^21^ LAMP primers were utilized to determine the infecting species. The MG-LAMP diagnosis was compared to results given by the local microscopist at the sites of study. Furthermore, the MG-LAMP data were compared to real-time PCR (PET-PCR)^22^-^24^ assays used as a reference test The feasibility and increased sensitivity of MG-LAMP compared to microscopy make this molecular diagnostic tool a good candidate to use in resource-limited communities, in areas where malaria transmission is low and active case detection is needed and to detect infection in patients with mixed infections and low parasite densities.

## MATERIALS AND METHODS

### Collection of clinical samples

This study was carried out between July and August 2017 in malaria outpatient clinics in three municipalities of Roraima, Brazil (Boa Vista, Pacaraima, and Rorainopolis). Written informed consent were obtained from all participants and blood was drawn by venipuncture. This study was approved by the Federal University of Roraima Ethical Committee (CAAE: 44055315.0.0000.5302).

All patients attending the outpatient’s health clinics for malaria screening were eligible to be enrolled in the study. Enrolled patients were tested for malaria by a trained local microscopist using 10% Giemsa-stained thick blood smear, and the diagnosis and parasitemia level were recorded for each patient. Additionally, all consenting patients filled out a clinical questionnaire. Information regarding whether the patient was symptomatic or not, their age and sex, and whether they had prior *Plasmodium* infections was documented. All of the sample processing, microscopy, DNA extraction and MG-LAMP assays were performed in Roraima.

### DNA Extraction

DNA was extracted from 200µL of whole blood using the QIAamp DNA Mini Kit (Qiagen). The DNA extraction protocol was slightly modified in that all of the spins were performed at 2,000g using a mini-centrifuge (Myfuge^™^) that was easily transported in the field setting as opposed to a centrifuge with adjustable speeds/time.

### LAMP method

To simplify the MG-LAMP procedure for ease-of-use in a simpler setting, a three-component ready-to-use kit was used: component I contained all the necessary reaction components for the assay (LAMP buffer: 40 mM Tris-HCL pH 8.8, 20 mM KCl, 16 mM MgSO_4_, 20 mM (NH_4_)SO_4_, 0.2% Tween-20, 0.8 M Betaine, and 2.8 mM of dNTPs and the primers); component II contained the Bst polymerase and component III contained 0.2% malachite green dye. To perform the assay, 13.8µL of Component I was mixed with 0.8µL of the Bst polymerase and 0.4uL of the malachite green dye for a final concentration of malachite green of 0.008%. This was mixed carefully and 5µL of DNA template was added. All samples were screened for *Plasmodium* using the genus assay as described previously^17, 19^ in a final reaction volume of 20µL. Samples were incubated for 1 hour at 63°C in a mini heat block (GeneMate, Bioexpress) to amplify the DNA. Following the 1-hour incubation, samples were removed from the heat block and allowed to cool for 15 minutes, the results were then scored by two independent readers as being positive (light blue/green) or negative (clear/colorless). A positive and negative control was included during each run using *P. falciparum* 3D7 DNA or nuclease free water, respectively.

*P. falciparum a*nd *P. vivax* species-specific MG-LAMP assays were carried out on all samples which were positive by the genus assay. These assays were performed in the similar way as the genus assays using the 3-component ready-to-use in-house kits prepared using previously published *P. falciparum* and *P. vivax* primers^20, 21^. Each reaction contained 5µL of isolated DNA in a final reaction volume of 20µL. Positive controls included *P. falciparum* sample and *P. vivax* positive sample. Nuclease free water was used for each assay as a negative control.

### PET-PCR method

DNA samples were brought back to the malaria branch laboratory at the CDC and a *Plasmodium* Genus-specific PET-PCR was performed, in duplicate, on all 91 samples as described previously with a few modifications^22^. The reactions were each 20µL containing 2X TaqMan Environmental Master Mix 2.0 (Applied Biosystems), 250nM of Genus forward Primer and FAM-Genus reverse primer, and 5µL of isolated DNA. The PET-PCR reaction was ran using an Agilent real-time PCR machine. The following cycling parameters were used: 15 minutes initial hot-start at 95°C followed by 45 cycles of denaturing at 95°C for 20 seconds, annealing at 63°C for 40 seconds, and an extension of 30 seconds at 72°C. A positive and negative control, 3D7 and nuclease free water respectively, were used during each run. Samples were designated as positive if they had a Ct value below 40.0 and negative if they had No Ct value or Ct values above 40.0.

Species-specific PET-PCR was performed, in duplicate, on all samples that were positive by the genus specific PET-PCR, using species-specific primers (Table 1). Two duplex reactions were set up to detect *P. ovale* together with *P. falciparum* and *P. malariae* together with *P. vivax.* The duplexed reactions were 20µL containing 2X TaqMan Environmental Master Mix 2.0 (Applied Biosystems), 250nM of FAM-*P. ovale* forward primer, 250nM *P. ovale* reverse primer, 250nM of *P. falciparum* forward primer, 125nM of HEX-*P. falciparum* reverse primer, 250nM *P. malariae* forward primer, 250nM FAM-*P. malariae*, 125nM *P. vivax* forward primer, 125nM HEX-*P. vivax* reverse primer and 5µL of isolated DNA. Reactions were ran using the same cycling conditions as the Genus PET-PCR. Positive controls consisting of samples with known *Plasmodium* species and nuclease free water as a negative control were included in every run. Samples were designated as positive if they had a Ct value below 40.0 and negative if they had No Ct value or Ct values above 40.0.

**Table 1:**
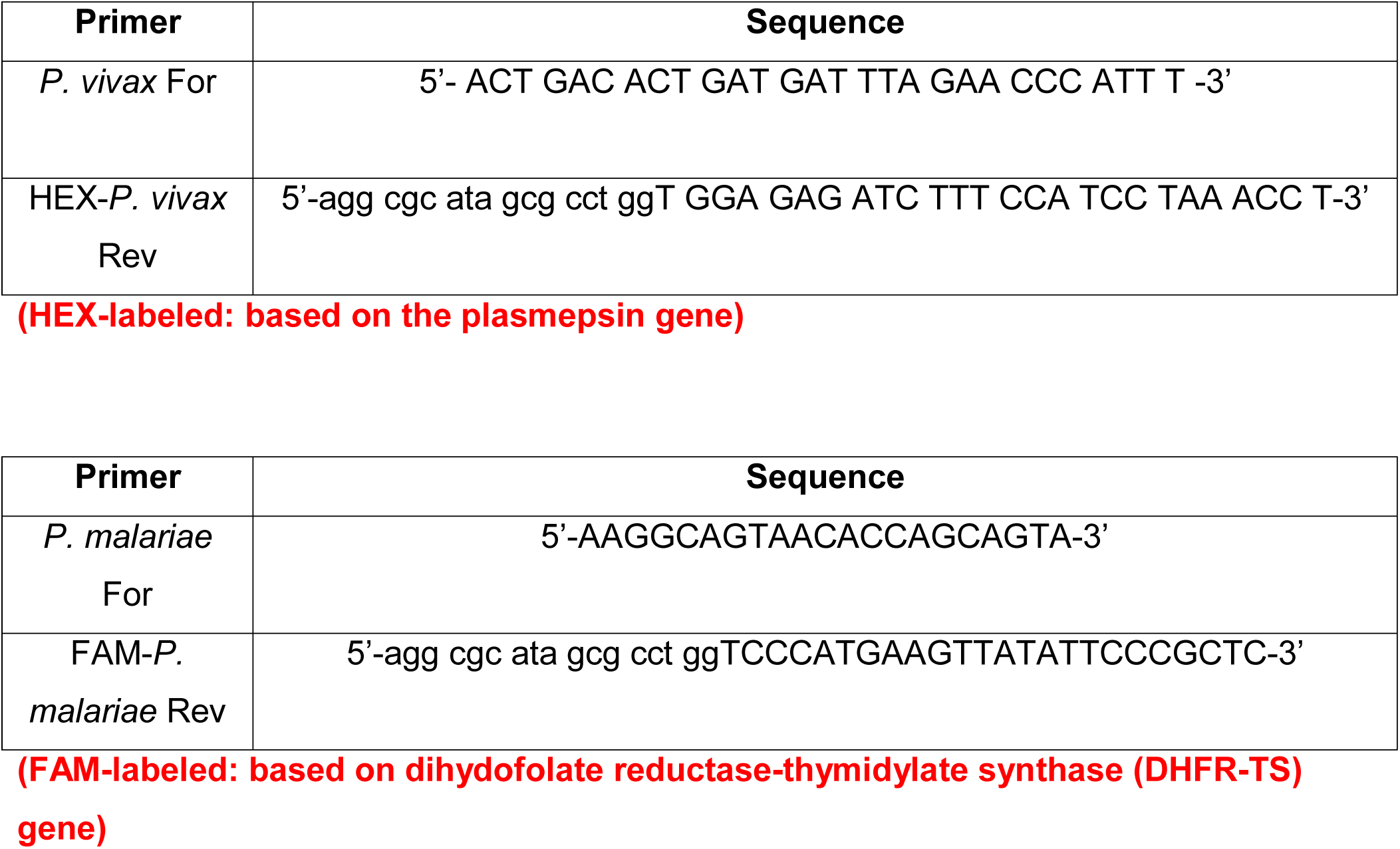
PET-PCR Primers utilized in the evaluation.

### Statistical analyses

Sensitivity and specificity tests were calculated as described previously^19^. The agreement between the human readers and diagnostic tests were assessed by calculating the kappa coefficients. 95% confidence intervals were calculated using MEDCALC^®^ and GraphPad.

## RESULTS

### Patient enrollment

A total of 91 patients presenting at the health clinics were enrolled in the study, Figure Sixty-five of these presented with malaria symptoms (axillary temperature ≥ 37.5 °C) and 26 presented no malaria symptoms. All the samples collected were tested for malaria parasites using microscopy, MG-LAMP and PET-PCR assays. Of the 91 enrolled patients, 86 (94.5%) reported to have had previous malaria infections while 4 (4.4%) had no previous infections and 1 (1.1%) did not provide this information.

**Figure 1:**
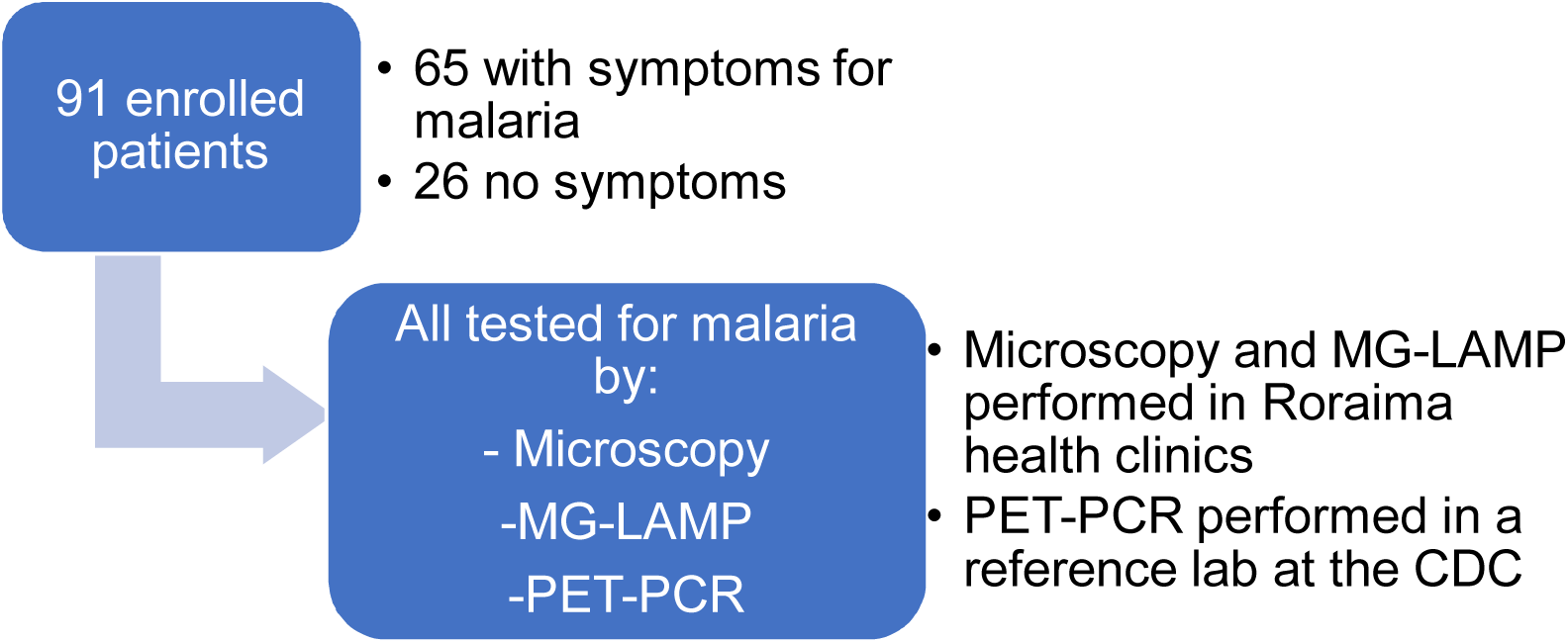
Summary of enrolled patients.

### Agreement between human readers for the MG-LAMP assay

Two independent readers were used to score the MG-LAMP tests to assess whether the sample was positive or negative. There was 100% agreement between the two readers (Kappa=1).

### Overall results of microscopy, MG-LAMP, and PET-PCR

A total of 91 samples were tested by microscopy, MG-LAMP and PET-PCR assay. Of the 91 samples, 33 (36%) were positive by microscopy, 39 (43%) were positive by MG- LAMP, and 40 (44%) were positive by PET-PCR. Species-specific reactions were carried out on all genus-positive samples using *P. falciparum* and *P. vivax* primers. A summary of the overall results obtained by each test are shown in Figure 2.

**Figure 2:**
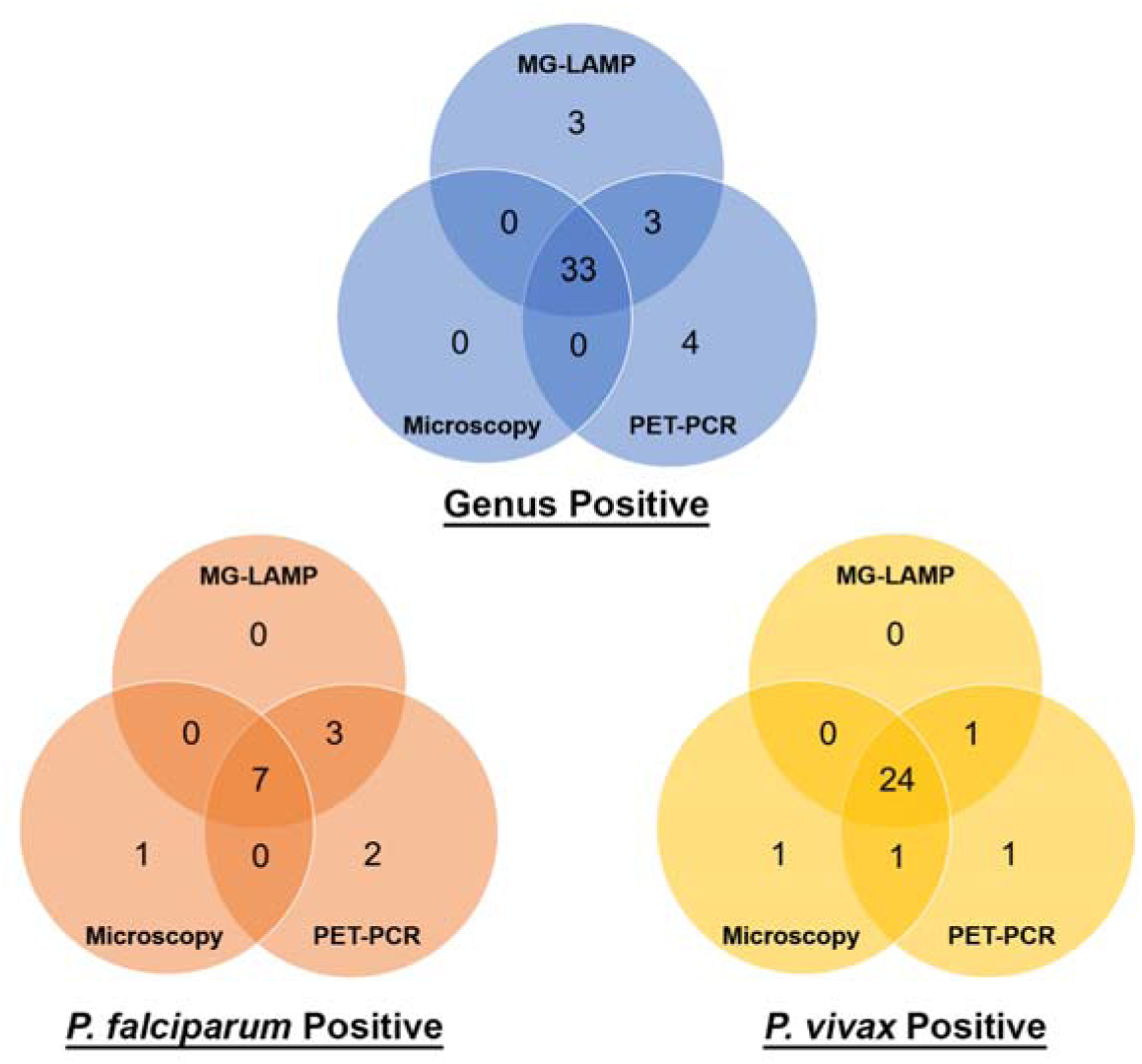
Summary of results for microscopy, MG-LAMP, and PET-PCR.

### Agreement of MG-LAMP to microscopy and PET-PCR

We observed that the MG-LAMP genus test and microscopy result (*Plasmodium* positive or negative) agreed 93.4% of the time (Kappa=0.86, 95% CI: 0.758 to 0.968). Furthermore, MG-LAMP analysis for *P. falciparum* and *P. vivax* diagnoses agreed with microscopy 96.7% (Kappa=0.81, 95% CI: 0.592 to 1.000) and 94.5% (Kappa=0.87, 95% CI: 0.754 to 0.980) of the time, respectively.

We found that *Plasmodium* genus assay, for MG-LAMP and PET-PCR, agreed 92.3% of the time (Kappa=0.84, 95% CI: 0.732 to 0.955). When comparing the *P. falciparum* and *P. vivax* MG-LAMP and PET-PCR assays we show a 97.8% (Kappa=0.89, 95% CI: 0.735 to 1.000) and 96.7% (Kappa=0.92, 95% CI: 0.839 to 1.000) agreement between the two tests, respectively. All samples were negative for *P. malariae* and *P. ovale* by microscopy and PET-PCR.

### Specificity and sensitivity of MG-LAMP and microscopy compared to PET-PCR

The sensitivity and specificity of the MG-LAMP assays and microscopy were calculated in comparison to PET-PCR used as a reference test, Table 2.

**Table 2:** Sensitivity and specificity of MG-LAMP and microscopy using PET-PCR as a reference.

|  | Method | Sensitivity | Specificity |
| --- | --- | --- | --- |
| **Genus** | **Microscopy** | 83% (95% CI:67.22%-92.66%) | 100% (95% CI: 93.02%-100.00%) |
|  | **MG-LAMP** | 90% (95% CI: 76.34%-97.21%) | 94% (95% CI: 83.76%-98.77%) |
| ***P. falciparum*** | **Microscopy** | 64% (95% CI: 93.02%-100.00%) | 99% (95% CI: 93.23%-99.97%) |
|  | **MG-Lamp** | 82% (95% CI: 48.22%-97.72%) | 100% (95% CI: 95.49%-100.00%) |
| ***P. vivax*** | **Microscopy** | 83% (95% CI: 65.28% - 94.36%) | 98% (95% CI: 91.20%- 99.96%) |
|  | **MG-LAMP** | 90% (95% CI: 73.47%- 97.89%) | 100% (95% CI: 94.13% - 100.00%) |

### Detection of mixed infections

Microscopy detected one mixed infection, here defined as infection with both *P. falciparum* and *P. vivax*. There were two mixed infections detected by MG-LAMP and six detected by PET-PCR (Table 3). In the four cases where the MG-LAMP did not detect the mixed infections identified by the PET-PCR, the Ct values were observed to be high, indicating low parasite density infections, Table 3.

**Table 3:** Detection of mixed infections by microscopy, MG-LAMP and PET-PCR. **^*^**Only one *P. vivax* parasite was seen under the microscopy

| **Sample** | **Microscopy diagnosis** | **MG-LAMP diagnosis** | **PET-PCR diagnosis** | **PET-PCR CT value for *P. falciparum*** | **PET-PCR CT value for *P. vivax*** |
| --- | --- | --- | --- | --- | --- |
| PC121 | *P. vivax* | Mixed | Mixed | 22.68 | 28.5 |
| PC123 | *P. falciparum* | Mixed | Mixed | 26.56 | 39.9 |
| BV237 | *P. vivax* | *P. vivax* | Mixed | 35.1 | 29.12 |
| BV217 | *P. vivax* | *P. vivax* | Mixed | 36.24 | 32.23 |
| BV239 | *P. vivax* | *P. falciparum* | Mixed | 31.37 | 35.38 |
| BV241 | *P. falciparum* | *P. falciparum* | Mixed | 29.43 | 39.92 |
| BV240 | Mixed* | *P. falciparum* | *P. falciparum* | 37.82 | No Ct |

### Detection of asymptomatic patients

Out of the twenty-six enrolled patients with no malaria symptoms, five were shown to be positive for malaria (asymptomatic)by the MG-LAMP and three by the PET-PCR assay. None of these were positive by microscopy (Table 4). Four of the five positive cases by MG-LAMP were only positive at the genus level and the infecting species could not be determined (Table 4).

**Table 4:** Asymptomatic patients detected by MG-LAMP and PET-PCR.

| Sample | Microscopy<br>Diagnosis | MG-LAMP<br>diagnosis | PET-PCR<br>Genus<br>(Ct value) | PET-PCR<br>*P. vivax*<br>(Ct value) | PET-PCR *P.*<br>*falciparum*<br>(Ct value) |
| --- | --- | --- | --- | --- | --- |
| RR09 | Negative | Genus only | Negative<br>(40.7) | Negative<br>(No Ct) | Negative<br>(No Ct) |
| RR10 | Negative | Genus only | Negative<br>(41.76) | Negative<br>(No Ct) | Negative<br>(No Ct) |
| RR37 | Negative | *P. vivax* | Positive<br>(32.74) | Positive<br>(35.96) | Negative<br>(No Ct) |
| RR41 | Negative | Genus only | Positive<br>(38.76) | Negative<br>(41.99) | Negative<br>(No Ct) |
| RR42 | Negative | Genus only | Negative<br>(40.74) | Negative<br>(41.69) | Negative<br>(No Ct) |
| RR53 | Negative | Negative | Positive<br>(34.99) | Positive<br>(39.09) | Negative<br>(No Ct) |

### Discordant Results

Seven samples were found to be discordant among the three tests (Table 5). Four of these samples were negative by microscopy and MG-LAMP but positive by PET-PCR. Three of these samples were only positive by PET-PCR genus test and negative by species tests, while one was positive by PET-PCR *P. vivax* (Table 5). In these four cases, the obtained Ct values by PET-PCR were all above 35.0 Three samples yielded a positive MG-LAMP genus test but were negative for the MG-LAMP *P. falciparum* and *P. vivax* tests and by both microscopy and PET-PCR (Table 5).

**Table 5:** Summary of discordant results.

| Sample | Microscopy diagnosis | MG-LAMP Genus diagnosis | PET-PCR Genus (Ct value) | PET-PCR *P. vivax* (Ct value) | PET-PCR *P. falciparum* (Ct value) |
| --- | --- | --- | --- | --- | --- |
| RR53 | Negative | Negative | Positive (34.99) | Positive (39.09) | Negative (No Ct) |
| BV235 | Negative | Negative | Positive (35.78) | Negative (No Ct) | Negative (No Ct) |
| RR01 | Negative | Negative | Positive (37.96) | Negative (No Ct) | Negative (No Ct) |
| BV236 | Negative | Negative | Positive (39.34) | Negative (No Ct) | Negative (No Ct) |
| RR09 | Negative | Positive | Negative (40.7) | Negative (No Ct) | Negative (No Ct) |
| RR10 | Negative | Positive | Negative (41.76) | Negative (No Ct) | Negative (No Ct) |
| RR42 | Negative | Positive | Negative (40.74) | Negative (41.69) | Negative (No Ct) |

## DISCUSSION

The findings presented in this study demonstrate the feasibility and accuracy of MG- LAMP as a malaria diagnostic test in a health clinic in a malaria endemic country. Importantly our data demonstrate that MG-LAMP is sensitive enough at identifying low-density infections and asymptomatic patients, which is important for malaria control and elimination efforts. Low parasitemia infections and asymptomatic cases are often missed by microscopy blood smear or standard RDT. In turn, these patients remain untreated and thus act as reservoirs for transmitting malaria. Additionally, we demonstrate that this assay, like the PET-PCR assay used as a reference test in this study, is capable of detecting mixed infections that microscopy missed. Treatment for malaria varies depending on the causative species. If a mixed infection goes undetected, the patient may not receive the appropriate medication, remaining ill and likely actively transmitting. Malaria elimination is incumbered by the lack of tools which are sensitive, portable, and easy to use. As a more sensitive and less subjective assay than microscopy and conventional RDTs, the MG-LAMP assay circumvents many of these issues, providing an idea alternative molecular tool for the detection of low-density infections. Previous studies have demonstrated that malaria LAMP assays in general are idea malaria diagnostics for the detection of malaria in asymptomatic patients^14^ and in the detection of non-falciparum infections.

Identification of asymptomatic patients and mixed infections is important for malaria control and elimination and is one of the advantages of having molecular tests. This appeared to be a benefit of both MG-LAMP and PET-PCR. The feasibility and sensitivity of MG-LAMP make this a great tool for screening large pools of asymptomatic patients in epidemiological surveillance studies. There were four instances where false negatives were observed using the MG-LAMP assay when using PET-PCR as a reference test (Table 5). In addition, four mixed infections were missed by the MG- LAMP that were detected by the PET-PCR assay. These missed infections were all shown to be of much lower parasite densities given the high Ct values (between 35 and 39) obtained in the PET-PCR assay. Extrapolation using our previously obtained PET-PCR data shows that a Ct value of 35.0 correspond to about 16 parasites/µL^16^; therefore, the missed samples had parasite densities of about 16 parasites/µL or below. These results imply that more sensitive primers for the detection of malaria using the MG-LAMP assay may be required. This would likely be improved by using LAMP primers with higher sensitivity in the future. Interestingly, there were three cases where LAMP yielded a positive genus result, while microscopy and PET-PCR were negative (Table 5). It is likely that these are false positives by the MG-LAMP assay. Alternatively, these patients may have true malaria infections not detected by the PET-PCR assay. Inconsistencies such as these are not uncommon especially when parasite densities are at or below the detection limits of a test. The reproducibility of PCR assays in the detection of samples with very low parasitemia was shown to alternate between positive and negative in about 38% of PCR replicates tested^25^. Similar inconsistencies, with low parasitemia samples, were observed in other studies^15, 26^. While PET-PCR remains a more sensitive assay overall when diagnosing low-density infections, it is a far more complicated procedure compared to the MG-LAMP, which requires costly equipment and resources. The performance of the MG-LAMP assay required only a small portable heat block and mini-centrifuge. The heat block used in this study had a 38-samples capacity allowing for screening of many samples at once. This aspect could be especially useful when screening large populations of people for malaria. However, a limitation of the current format of the MG-LAMP is the fact that the LAMP buffer and polymerase utilized required a cold chain, which is not ideal in more resource-limited settings. In addition, the necessity to isolate DNA using a blood kit was time consuming and expensive, however previous publications demonstrated the compatibility of a boil- and-spin DNA isolation method with MG-LAMP^19^. The use of boil-and-spin DNA isolation could be further explored in future field studies. Therefore, improvements to make the MG-LAMP assay cold-chain free will be required, if this tool will be used in future epidemiological surveillance studies.

Overall, MG-LAMP provides a portable, user-friendly method for diagnosing malaria, and it is less subjective and more sensitive than microscopy. Importantly, MG-LAMP has the capacity to test at least 38 samples at a time allowing for the screening of large number of samples which is appealing when large scale studies are necessary e.g. in community surveillance studies. The current MG-LAMP assay was limited in its ability to detect mixed infection and extremely low-density infections, but otherwise proved to be an advantageous tool for diagnosing malaria in the field and opens news perspectives in the implementation of surveillance in malaria elimination campaigns.

## ACKNOWLEDGMENTS

We would like to thank all the patients who enrolled in this study and made this work possible and the Secretary of Health of Roraima and the local malaria control program team of Boa Vista, Pacaraima and Rorainopolis for their logistical support This work was supported by the US National Institutes of Health to H.M.K (T32AI060546). JOF is recipient of Research Productivity Fellowships from the Conselho Nacional de Desenvolvimento Científico e Tecnológico (CNPq).

